# Associations between Fractional Anisotropy in White Matter Bundles and AT(N) Plasma Biomarkers in Individuals with Cognitive Complaints without dementia

**DOI:** 10.64898/2026.11.25.768865

**Authors:** Patricio F. Riquelme, Isaías Mery, Sebastián Aguilera, Fernando Henríquez, Vicente Medel, Pamela C L. Ferreira, Bruna Bellaver, Thomas K. Karikari, Paulina Orellana, Teresita Ramos, Tharick A. Pascoal, Daniela Thumala, Cecilia Okuma, Patricia Lillo, J. César Cardenas, Christian González-Billault, Felipe A. Court, Claudia Duran-Aniotz, Pamela Guevara, Andrea Slachevsky

## Abstract

Age-related cognitive decline is preceded by subtle neurochemical and structural brain alterations that remain insufficiently characterized, particularly in individuals with cognitive complaints but without dementia and in populations from the Global South. AT(N) plasma biomarkers reflecting amyloid pathology, tau phosphorylation, astroglial reactivity, and axonal injury have emerged as accessible indicators of early neurodegenerative processes; yet their relationship with white matter microstructural integrity and the extent to which this relationship is confounded by co-occurring cerebrovascular pathology remain unclear. We investigated associations between AT(N) plasma biomarkers, including the Aβ42/Aβ40 ratio, phosphorylated tau-217 (p-tau217), glial fibrillary acidic protein (GFAP), and neurofilament light chain (NfL), and white matter fractional anisotropy (FA) derived from diffusion MRI in 135 community-dwelling Chilean older adults with cognitive complaints (GERO cohort). Whole-brain correlational tractography was performed controlling for age and education, and a secondary analysis (n = 124) additionally adjusted for intracranial volume-normalized white matter hyperintensity (WMH) burden after excluding individuals with Fazekas scores >2. Implicated tracts were anatomically labeled using the HCP-1065 atlas. In the primary analysis, higher GFAP and p-tau217 were inversely associated with FA across periventricular tracts, including the corpus callosum tapetum, fornix, optic radiations, and posterior thalamic radiations; NfL showed a similar but predominantly callosal pattern; and direct associations were detected in limbic bundles such as the cingulum and fornix. After WMH adjustment, GFAP and NfL associations were preserved and spatially amplified into projection and association tracts, whereas p-tau217 associations did not survive adjustment, and an inverse Aβ42–FA association in the fornix persisted. These findings suggest that plasma GFAP and NfL index white matter microstructural processes biologically distinct from macroscopic small-vessel disease, whereas p-tau217 captures FA variance that is topographically inseparable from co-localized vascular pathology, at least in moderate-sized cross-sectional designs. Together, these results advance understanding of the neuroanatomical correlates of blood-based biomarkers at pre-dementia stages in Latin American populations.

## Introduction

Age-related cognitive impairment and dementia represent rapidly growing public health challenges worldwide, driven by the accelerated aging of the global population (Suemoto et al., 2026). Neurocognitive disorders, including Alzheimer’s disease dementia (ADD), are characterized by a prolonged preclinical phase that may extend for 15–20 years prior to the onset of overt dementia (Serrano-Pozo et al., 2026). Identifying individuals during these early stages is critical, as it provides a window of opportunity for preventive strategies and disease-modifying interventions. Evidence from high-income countries indicates that molecular and neuroimaging biomarkers can contribute substantially to identifying individuals at risk during preclinical phases, or those presenting with cognitive complaints in the absence of dementia (Shaikh et al., 2025).

Within this population, neuroimaging markers such as cortical atrophy, white matter hyperintensities (WMH), and altered functional connectivity have been consistently reported relative to cognitively unimpaired (CU) individuals, even in the absence of objective neuropsychological impairment (Dhana et al., 2022; Pievani et al., 2024; Ribaldi et al., 2024). These findings suggest that neurobiological changes may precede measurable cognitive decline, highlighting the relevance of sensitive imaging biomarkers for early disease detection.

Among these markers, white matter microstructural integrity has emerged as a particularly relevant neuroimaging biomarker, as alterations in white matter are thought to occur at very early stages of cognitive impairment (Chao et al., 2022; Wang et al., 2012; Wei et al., 2021). Fractional anisotropy (FA), derived from diffusion tensor imaging (DTI), is widely used to estimate the microstructural properties of white matter tracts (Meneguzzo et al., 2019). In individuals with mild cognitive impairment (MCI) and ADD, reduced FA in major tracts, including the corpus callosum, fornix, and parahippocampal cingulum, has been associated with cognitive dysfunction, myelin degeneration, and axonal damage (Bergamino et al., 2021; Kantarci, 2014). Importantly, these microstructural changes may precede macroscopic gray matter atrophy and white matter lesions such as WMHs, positioning FA as a sensitive biomarker for early disease processes (Kantarci, 2014; Li et al., 2023; Sun et al., 2025; Teipel et al., 2014). Furthermore, FA alterations have been shown to correlate with amyloid and tau positron emission tomography (PET), which are currently considered the gold-standard methods for characterizing Alzheimer’s pathology along the ADD continuum (Carlson et al., 2021; Sintini et al., 2019). However, PET imaging remains invasive, costly, and largely inaccessible in the Global South (Daly et al., 2026), underscoring the need for more accessible biomarkers that reflect early structural brain changes.

In this context, AT(N) plasma biomarkers have gained increasing attention as minimally invasive, cost-effective tools with high diagnostic precision for detecting neurodegenerative processes, even at early stages (Caviedes et al., 2026; Duran-Aniotz et al., 2026; Pais et al., 2023; Palmqvist et al., 2020). Neurofilament light chain (NfL) is a well-validated marker of axonal injury, and elevated plasma levels have been shown to predict accelerated cognitive decline and increased brain atrophy in individuals with cognitive complaints (Bangen et al., 2021; Götze et al., 2024). Glial fibrillary acidic protein (GFAP), a marker of reactive astrogliosis, has been associated with cerebral amyloid pathology, with elevated plasma levels observed even in individuals with subjective cognitive complaints (Chatterjee et al., 2021, 2022; Pereira et al., 2021). Among tau biomarkers, phosphorylated tau at threonine 217 (p-tau217) has demonstrated particularly high specificity for ADD pathology, differentiating ADD from other neurodegenerative diseases well before the onset of clinical impairment (Algeciras-Schimnich et al., 2026; Caviedes et al., 2026; Giacomucci et al., 2025; Orellana et al., 2026; Palmqvist et al., 2020). Finally, the plasma Aβ42/Aβ40 ratio serves as an indirect indicator of cerebral amyloid burden, with lower ratios reflecting increased amyloid deposition. In individuals with cognitive complaints, reductions in this ratio correlate with amyloid PET findings and improve diagnostic accuracy when combined with GFAP (Algeciras-Schimnich et al., 2026; Chatterjee et al., 2022; Kang et al., 2025).

Despite the growing evidence supporting the utility of AT(N) plasma biomarkers for early ADD detection, their relationship with early indicators of brain damage, particularly white matter microstructural integrity, remains insufficiently characterized. Clarifying these associations is essential for strengthening the biological validity of plasma biomarkers as surrogates for early neurodegenerative and glial-related structural changes in the aging brain.

In a previous study conducted by our group, we demonstrated that WMH volume was associated with cardiovascular risk factors, particularly diastolic blood pressure, but not with plasma biomarkers of neurodegeneration (Riquelme-Contreras et al., 2026). As WMHs represent a relatively late, macroscopic manifestation of white matter damage, these findings suggest that vascular mechanisms may precede, or operate independently of, detectable neurodegenerative biomarker changes during early cognitive decline. Accordingly, assessing white matter microstructure using diffusion-derived metrics, such as FA, could capture subtler and earlier alterations that are not yet expressed as WMHs, thereby providing a more sensitive window into the initial neurobiological processes linking white matter vulnerability to plasma markers of neurodegeneration and glial reactivity.

Therefore, in the present study, we aimed to investigate the associations between AT(N) plasma biomarkers and the integrity of white matter fiber bundles, as measured by FA derived from DTI. Further, we explored the influence of WMH volume on these associations. This study sought to characterize these associations in the GERO cohort, a unique community-based sample of Chilean older adults with low educational attainment and cognitive complaints without dementia, encompassing a spectrum ranging from subjective cognitive complaints without objective impairment to objective cognitive impairment, in whom extensive clinical, cognitive, neuroimaging, and biomarker assessments have been performed (Slachevsky et al., 2020). While this cohort parallels well-established community-based studies from North America and Europe, it provides novel and much-needed evidence from a Latin American context, contributing to a more globally representative understanding of early neurobiological changes associated with age-related cognitive decline. We hypothesize that AT(N) plasma biomarkers will be associated with microstructural changes in white matter fiber bundles, with variable influence from WMH burden, reflecting differential relationships between cerebral small vessel disease and neurodegeneration, reactive astrogliosis, and axonal injury.

## Material and methods

### Participants

We included 135 individuals (108 females) selected from a Chilean community-based cohort of individuals with cognitive complaints without dementia (GERO Cohort), ranging from those with subjective cognitive complaints and no objective cognitive impairment to those with objective cognitive impairment (Slachevsky et al., 2020). In brief, we employed a door-to-door recruitment strategy to identify and enroll community-dwelling adults aged 70 years and older from the general population. Household visits were conducted across three districts of Santiago de Chile, which were selected for their socioeconomic diversity to ensure representativeness. Participants were included if they reported cognitive complaints themselves or through a reliable proxy, scored 0.5 or less on the Clinical Dementia Rating Scale (CDR), lived in the community, and had a knowledgeable informant. For self-reported cognitive complaints, the questionnaire included 11 items from Rabin et al. (2015), along with additional questions developed and preliminarily validated by our group. Proxy-reported cognitive complaints were assessed using the AD8 questionnaire and other items validated by our group (Galvin et al., 2005; Molina-Donoso, Parrao, et al., 2023; Muñoz et al., 2010; Parrao et al., 2018; Rabin et al., 2015; Slachevsky et al., 2017). Participants were included if they responded positively to a predefined set of questions (see Supplementary Material S1). We excluded individuals with dementia, as identified by screening tests that documented the concurrent presence of cognitive impairment (scores < 21 on the Mini-Mental State Examination, MMSE) and functional impairment (scores > 2 on the Pfeffer Functional Activities Questionnaire, PFAQ) on screening assessment (Arevalo-Rodriguez et al., 2015; Molina-Donoso, González-Hernández, et al., 2023; Pfeffer et al., 1982). Participants were also excluded from the study if they had a history of institutionalization, illiteracy, visual and auditory deficits impeding cognitive evaluation and/or the capacity to maintain independent living, severe mobility impairment, severe psychiatric or neurological disorders, or a lethal disease with a prognosis of less than one year of survival. Additionally, a trained neurologist evaluated all participants to verify that they met the inclusion criteria.

Hand dominance was assessed during the cohort’s sociodemographic interview. In the initial sample (n = 135), 97.8% of participants were right-handed.

Additionally, the following data were available for all selected individuals: a multidimensional neuropsychological assessment covering episodic memory (Free and Cued Selective Reminding Test [FCSRT]), attention and executive function (Color Trails Test Parts A and B [CTT-A/B]), and language (Sydney Language Battery Naming subtest [SydBat Naming]) (for details, see Slachevsky et al., 2020); diffusion magnetic resonance imaging (dMRI); the neurodegeneration blood biomarkers Aβ42 and Aβ40; the Aβ42/Aβ40 ratio derived from these markers; and p-tau217. Based on evidence demonstrating (1) the efficacy of p-tau217 and the Aβ42/Aβ40 ratio in detecting amyloid pathology (Ashton et al., 2024; Mielke et al., 2022; Palmqvist et al., 2024), (2) the capacity of p-tau217 to differentiate ADD patients from healthy controls using data from this cohort (Orellana et al., 2026), and (3) its capacity to differentiate between clinical groups (Caviedes et al., 2026), we used both plasma biomarkers as measures of neurodegeneration.

Regarding glial markers, glial fibrillary acidic protein (GFAP) and neurofilament light chain (NfL) were selected for their reported performance in detecting astrogliosis and axonal damage, respectively, and their correlation with neurodegeneration, amyloid status, and preclinical stages of dementia (Caviedes et al., 2026; Chatterjee et al., 2023; Efe et al., 2026; Ingannato et al., 2024; Orellana et al., 2026; Pereira et al., 2021).

To obtain these biomarkers, blood samples were collected at the subject’s home using a vacuum system with venous access via 21 G or 23 G needles, depending on the subject’s venous caliber. Specifically, 12 mL of blood was extracted, yielding 4 mL of plasma. However, the biomarkers studied were analyzed using 1 mL of plasma, equivalent to 3 mL of blood. Once extracted, samples were divided into aliquots and frozen at -80°C. Samples were shipped to the University of Pittsburgh (Pennsylvania, USA) under continuous temperature monitoring to maintain the cold chain during transport until laboratory processing. Plasma biomarkers Aβ42, Aβ40, p-tau217, GFAP, and NfL were quantified using a Neurology 4-Plex E (#103670) commercial assay kit (Quanterix, Billerica, MA, USA). More details about the sample extraction and processing can be found in our previous work (Orellana et al., 2026). More details about the recruitment process and a complete description of the study protocol are available elsewhere (Slachevsky et al., 2020). This published study protocol details the sample size calculation and justification.

The GERO project was approved on May 5, 2015, by the Scientific Ethics Committee of the Eastern Metropolitan Health Service of Santiago, Chile. No approval number was assigned. All participants provided written informed consent in accordance with the principles of the Declaration of Helsinki, as approved by the same ethics committee. The study protocol is registered at ClinicalTrials.gov (NCT04265482), and recruitment was conducted between May 2017 and July 2021.

### MRI acquisition and processing

Diffusion MRI images were acquired using a 3T Skyra scanner (Siemens) at the Neurosurgery Institute Dr. Alfonso Asenjo (Santiago, Chile), with the following acquisition parameters: single-shell acquisition, repetition time (TR) = 10,000 ms, echo time (TE) = 92 ms, field of view (FoV) = 240 mm, voxel size = 2.0 × 2.0 × 2.0 mm, slice thickness = 2 mm, b-value = 1000 s/mm^2^, and 30 diffusion-encoding directions.

Although multi-shell acquisitions enable more complex diffusion models, single-shell DTI with 30 gradient directions at b = 1000 s/mm^2^ has been shown to provide reliable FA estimates for tractography-based analyses in aging cohorts and is consistent with widely validated protocols for large-scale white matter microstructure studies (Bergamino et al., 2021; Teipel et al., 2014). To obtain DTI metrics, images were converted to NIFTI format (.nii) and analyzed using DSI Studio (https://dsi-studio.labsolver.org/). Preprocessing considered the diffusion gradient intensity (.bval) and diffusion direction vectors (.bvec) to generate .src files, followed by visual quality control to ensure image integrity (F.-C. Yeh, Zaydan, et al., 2019). Additionally, a brain mask was generated to remove background noise and improve reconstruction efficiency. Next, eddy current and head-eye motion correction were applied, followed by computation of the diffusion local model using q-space diffeomorphic reconstruction (QSDR). QSDR reconstruction parameters included an ODF sampling count of 1500, a diffusion sampling length ratio of 1.25, and normalization to the ICBM-152 template (F.-C. Yeh & Tseng, 2011). DTI-derived metrics, including FA, were then calculated. Finally, a connectometry database for FA was created using the .fib files from all participants (F.-C. Yeh et al., 2016).

### Statistical analysis

Descriptive statistics were used to summarize sex, age (years), education (years), neuropsychological and clinical characteristics, available blood biomarkers, and TIV-normalized WMH lesion load, including mean, standard deviation (SD), minimum, and maximum values. Correlational tractography (F.-C. Yeh et al., 2021) was performed to identify white matter pathways in which FA was significantly correlated with each plasma biomarker (Aβ42/Aβ40, p-tau217, GFAP, and NfL). A nonparametric Spearman partial correlation was used to assess these associations, with the effects of age and years of formal education removed via multiple regression. The statistical significance of the correlation was examined using a permutation test. A T-score threshold of 2.5 was applied in the fiber-tracking algorithm (F.-C. Yeh et al., 2013). A whole-brain seeding region was employed (MNI coordinates: 39, 49, 29). The resulting tracts were filtered using topology-informed pruning (F.-C. Yeh, Panesar, et al., 2019) with 16 iterations. A minimum length threshold of 15 voxels was used to select tracks. To estimate the false discovery rate (FDR), 4000 randomized permutations were applied to the group labels to obtain the null distribution of the track length (FDR < 0.05).

Further, tracts showing significant correlations were anatomically identified using probabilistic maps based on the HCP-1065 white matter tract atlas (F. C. Yeh, 2022), which includes 64 major white matter pathways. This anatomical identification step was performed using Python (version 3.10.14), along with the NumPy (Harris et al., 2020) and NiBabel v. 5.2.1 (Brett et al., 2024)

To address the potential confounding role of cerebrovascular white matter damage on DTI-derived metrics, and given that several implicated tracts traverse periventricular regions where WMH and microstructural FA changes are known to spatially overlap, we conducted an additional analysis incorporating WMH burden into the connectometry model. For this analysis, participants with Fazekas scores >2 in either the periventricular or deep white matter regions were excluded (n = 11), yielding a subsample of 124 individuals. Total WMH volume, normalized by intracranial volume (TIV), was then included as an additional covariate in the connectometry model, alongside age and years of formal education. This approach allowed us to evaluate whether the associations between plasma biomarkers and FA observed in the primary analysis were independent of co-occurring macroscopic cerebrovascular pathology.

The WMH data were obtained by automated segmentation of FLAIR MRI images. The acquisition parameters, preprocessing, and segmentation details can be found in our previous work (Riquelme-Contreras et al., 2026).

To quantify the strength of the associations, fractional anisotropy (FA) was extracted for each participant from each significant cluster of tracts, identified by correlational tractography (FDR < 0.05; positive and negative associations separately), using DSI Studio. Spearman partial correlations were then computed in R (v4.3.3, *ppcor* package) between tract-averaged FA and the corresponding plasma biomarker. These analyses used the same covariates and participant subsamples as the corresponding connectometry model: age and education, plus total white matter lesion volume in the WMH-adjusted model. Because tracts clusters were selected based on their association with each biomarker, these coefficients are reported as descriptive effect-size estimate rather than independent tests. The analysis scripts can be consulted in Supplementary Material S2.

Finally, as a sensitivity analysis, we repeated the analytical pipeline in participants classified as having subjective cognitive complaints (SCC) without objective cognitive impairment, based on performance in global cognition (MoCA) and episodic memory (FCSRT) using locally validated normative scores, according to the procedure previously described by our group (Henríquez et al., 2026).

## Results

### Demographics, clinical, and biomarkers

The average age of the participants was 76.09 years (SD = 4.93), with an average of 9.36 years of formal education (SD = 4.66). Furthermore, the sample exhibited a mean total CDR score of 0.45 (SD = 0.44) and a mean MMSE score of 27.09 (SD = 2.28). Regarding domain-level neuropsychological performance, several participants scored below conventional cut-offs on individual measures across cognitive domains (Table 1). However, isolated low performance was not used to classify participants as having dementia; classification was based on the overall cognitive and functional profile, including functional independence as assessed with the PFAQ (Forno et al., 2025).

**Table 1:** Description of demographic, clinical data, and plasma biomarker levels in the sample.

| **Variable** | **Mean (Median)** | **SD** | **Min** | **Max** |
|---|---|---|---|---|
| *Sociodemographics* |  |  |  |  |
| Sex (M : F), % = 27 : 108, 20 : 80 % |  |  |  |  |
| Age (years) | 76.09 (75) | 4.93 | 69 | 92 |
| Education (years) | 9.36 (8) | 4.66 | 0 | 24 |
| *Clinical and global cognition* |  |  |  |  |
| MMSE Total Score | 27.09 (28) | 2.28 | 20 | 30 |
| MoCA Total Score | 21.53 (22) | 4.38 | 11 | 29 |
| CDR Total Score | 0.45 (0.5) | 0.16 | 0 | 0.5 |
| PFAQ Total Score | 0.76 (0) | 1.28 | 0 | 8 |
| *Multi-domain neuropsychological assessment* |  |  |  |  |
| Semantic memory - FCSRT total recall | 47.31 (48) | 1.62 | 34 | 48 |
| Episodic memory - FCSRT differed free recall | 11.67 (12) | 2.66 | 3 | 16 |
| Attention - CTT A (sec) | 98.44 (83) | 53.83 | 24 | 360 |
| Executive function - CTT B (sec) | 234.12 (194) | 134.16 | 59 | 540 |
| Language – SydBat Naming | 24.74 (25) | 3.27 | 13 | 30 |
| *Plasma biomarkers and WMH burden* |  |  |  |  |
| Aβ42 (pg/ml) | 6.48 (6.40) | 1.773 | 1.376 | 11.810 |
| Aβ40 (pg/ml) | 108.96 (105.87) | 21.29 | 58.749 | 196.607 |
| Aβ42/Aβ40 ratio | 0.05 (0.060) | 0.01 | 0.015 | 0.092 |
| p-tau217 (pg/ml) | 0.23 (0.167) | 0.20 | 0.025 | 1.728 |
| NfL (pg/ml) | 29.91 (27.88) | 14.01 | 10.131 | 93.606 |
| GFAP (pg/ml) | 138.50 (124.79) | 70.80 | 36.982 | 503.720 |
| Total WMH (TIV normalized) | 0.00 (0.008) | 0.008 | 8.16*10^-4 | 0.051 |

The average plasma concentrations of Aβ42 and Aβ40 were 6.48 pg/mL (SD = 1.77) and 108.96 pg/mL (SD = 24.29), respectively. For p-tau217, the mean value was 0.23 pg/mL (SD = 0.20); GFAP averaged 138.50 pg/mL (SD = 70.80); and NfL averaged 29.91 pg/mL (SD = 14.01). Boxplots showing the distribution of AT(N) plasma biomarkers in the sample, along with the complete biomarker values, are provided in the Supplementary Material S3.

Demographic, clinical, and summary statistics for all blood biomarkers are presented in Table 1.

### Associations of FA with Aβ42/Aβ40

Correlational tractography exploring direct associations between Aβ42/Aβ40 plasma levels and FA revealed a particularly pronounced involvement of periventricular fibers, including the corpus callosum tapetum (4.35%), right optic radiation (1.96%), right occipital corticopontine tract (1.79%), and right posterior thalamic radiation (1.72%). The full list of identified tracts is shown in Table 2, with a graphical representation provided in Figure 1.

**Table 2:** Results of the anatomical bundle identification regarding direct associations between FA and Aβ42/Aβ40 plasmatic levels (FDR < 0.05, n=117). **Legend**: Bundle voxel n: number of voxels for the bundle according to the HCP 1065 white matter atlas; Bundle voxel int: number of fibers with intersection between the atlas and fibers correlated; %: percent of the total bundle compromised by the association.

| **Bundle** | **Bundle voxel n** | **Bundle voxel int** | **%** |
|---|---|---|---|
| Corpus callosum tapetum | 44230 | 1925 | 4.352 |
| Right optic radiation | 21010 | 413 | 1.965 |
| Right corticopontine tract occipital | 26345 | 473 | 1.795 |
| Right thalamic radiation posterior | 35747 | 615 | 1.72 |
| Corpus callosum forceps major | 117410 | 1874 | 1.596 |
| Right corticostriatal tract posterior | 51564 | 585 | 1.134 |
| **Spearman rho (all tracts) = 0.225 (p = 0.015), IC [2.5 , 97.5] = 0.049 , 0.397. Adjusted β = 0.229** |  |  |  |

**Figure 1:**
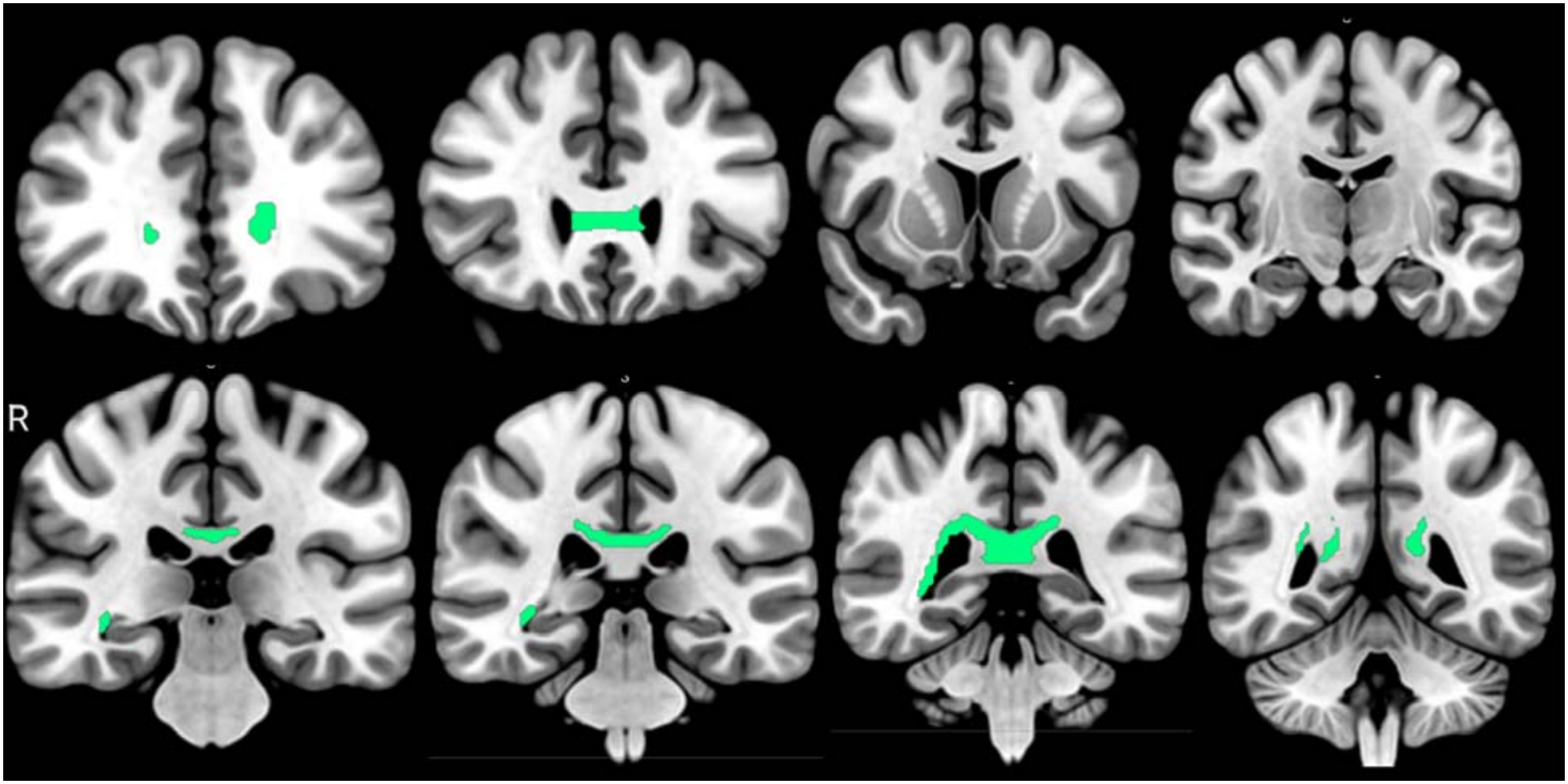
White matter tracts showing direct associations between FA and plasma Aβ42/Aβ40 levels (correlational tractography; Spearman partial correlation controlling for age and education; T-score threshold = 2.5; FDR < 0.05). Corpus callosum tapetum and right optic radiation show the highest proportions of compromised fibers (4.35% and 1.97%, respectively). Colors represent tract identity; numbers indicate the percentage of tract volume with significant FA-biomarker association relative to the HCP-1065 atlas definition.

Regarding the inverse associations between FA and Aβ42/Aβ40 plasma levels, the correlation coefficient was not significant (Spearman rho = -0.158, p = 0.088, IC 2.5 – 97.5% = -0.350, 0.030). See supplementary material S2 to explore the complete coefficients table.

### Associations of FA with p-tau217

Inverse associations between p-tau217 and FA mainly affected bundles within the posterior periventricular white matter, with the strongest associations observed in the corpus callosum tapetum (8.86%), right fornix (3.38%), and left optic radiation (3.29%). The full list of anatomically identified bundles is shown in Table 3, with an anatomical map of these associations represented in Figure 2.

**Table 3:** Results of the anatomical bundle identification regarding inverse associations between FA and p-tau217 plasmatic levels (FDR < 0.05, n=126). For a description of the table headings, see Table 2.

| **Bundle** | **Bundle voxel n** | **Bundle voxel int** | **%** |
|---|---|---|---|
| Corpus callosum tapetum | 44230 | 3919 | 8.860 |
| Right fornix | 2896 | 98 | 3.383 |
| Left optic radiation | 23529 | 775 | 3.293 |
| Corpus callosum forceps major | 117410 | 2287 | 1.947 |
| Left fornix | 2397 | 39 | 1.627 |
| Left thalamic radiation posterior | 38864 | 462 | 1.188 |
| Right thalamic radiation posterior | 35547 | 374 | 1.046 |
| Left corticostriatal tract posterior | 51578 | 534 | 1.035 |
| Left corticopontine tract occipital | 26840 | 273 | 1.017 |
| Right optic radiation | 21010 | 211 | 1.004 |
| **Spearman rho (all tracts) = -0.288 (p = 0.001), IC [2.5 , 97.5] = -0.430 , -0.131. Adjusted β = -0.267** |  |  |  |

**Table 4:** Results of the anatomical bundle identification regarding inverse associations between FA and GFAP plasmatic levels (FDR < 0.05, n=117). For a description of the table headings, see Table 2. See Supplementary Material for the complete report of the identified bundles.

| Bundle | Bundle voxel n | Bundle voxel int | % |
|---|---|---|---|
| Corpus callosum tapetum | 44230 | 5424 | 12.263 |
| Left fornix | 2397 | 124 | 5.173 |
| Left thalamic radiation posterior | 38864 | 1940 | 4.991 |
| Left optic radiation | 23529 | 791 | 3.361 |
| Right optic radiation | 21010 | 689 | 3.279 |
| Corpus callosum forceps major | 117410 | 3471 | 2.956 |
| **Spearman rho (all tracts) = -0.274 (p = 0.002), IC [2.5 – 97.5] = -0.441, -0.097. Adjusted β = -0.225** |  |  |  |

**Figure 2:**
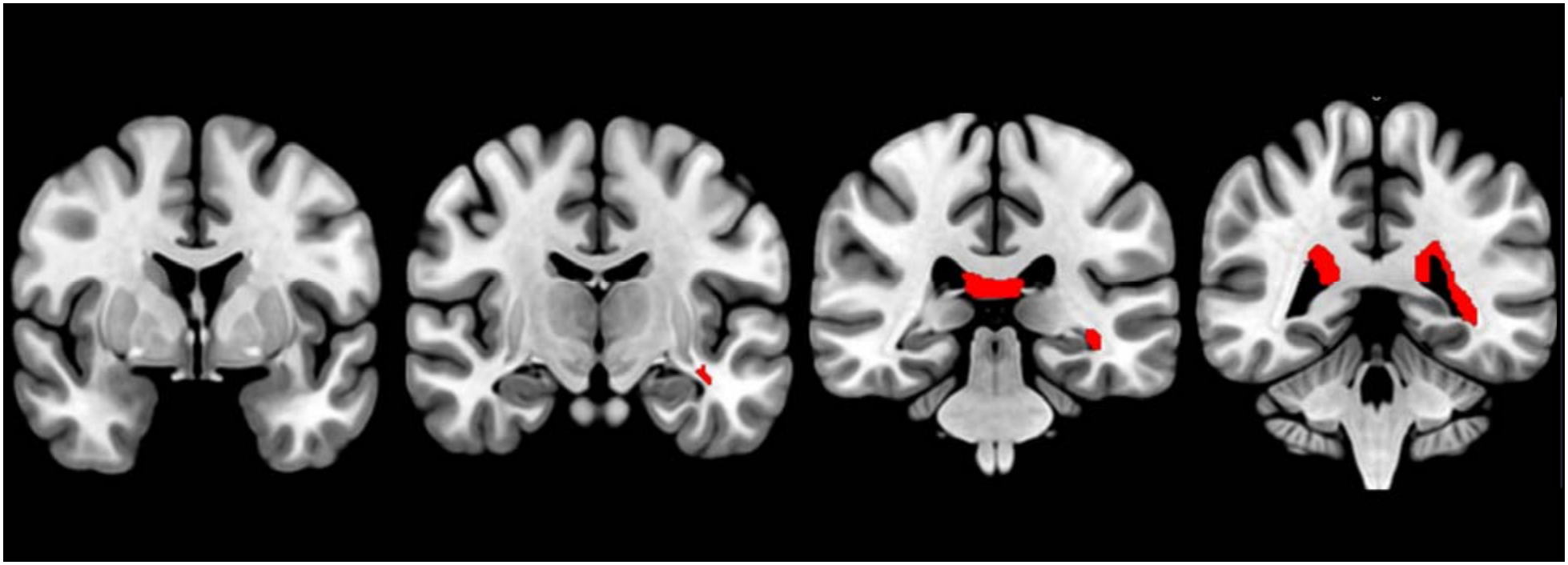
White matter tracts showing inverse associations between FA and plasma p-tau217 levels (correlational tractography; Spearman partial correlation controlling for age and education; T-score threshold = 2.5; FDR < 0.05). Corpus callosum tapetum (8.86%), right fornix (3.38%), and left optic radiation (3.29%) show the highest proportions of compromised fibers. The distribution is predominantly periventricular and posterior.

**Figure 3:**
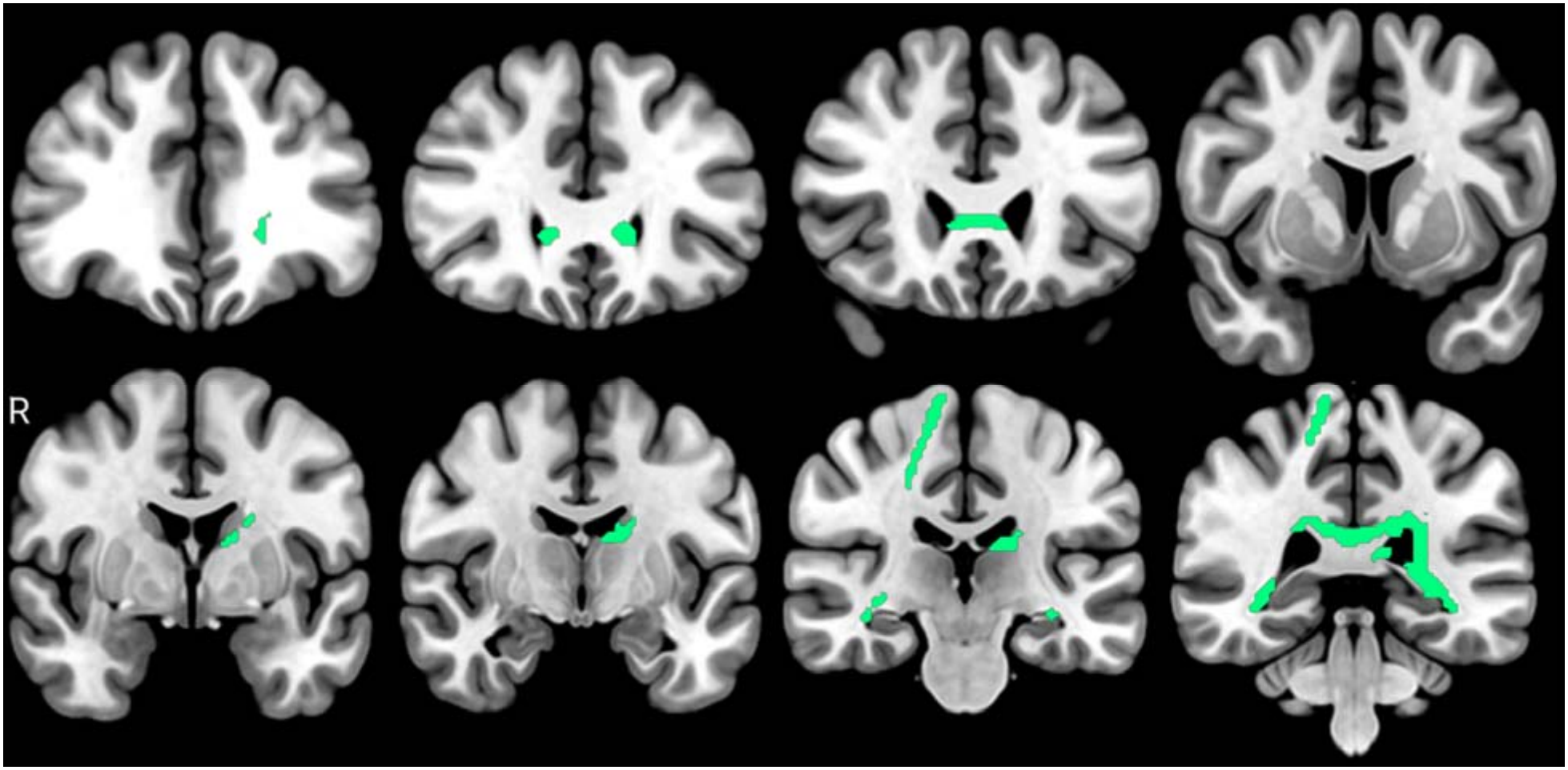
White matter tracts showing inverse associations between FA and plasma GFAP levels (correlational tractography; Spearman partial correlation controlling for age and education; T-score threshold = 2.5; FDR < 0.05). Corpus callosum tapetum (12.26%), left fornix (5.17%), and bilateral posterior thalamic radiations show the highest proportions of compromised fibers. The complete list of identified tracts is provided in Table 5 and Supplementary Material.

Regarding direct associations, although bundles within the limbic circuit exhibited associations, once Spearman correlation coefficient was calculated, it did not reach significance (Spearman rho = 0.174, p = 0.051, IC 2.5 – 97.5% = -0.008, 0.347). See supplementary material to explore the complete coefficients table.

### Associations of FA with GFAP

When calculating inverse associations between GFAP and FA, we found the highest number of bundles showing significant associations, with pronounced involvement of the posterior periventricular white matter. The corpus callosum tapetum had the highest percentage of associated fibers (12.26%), followed by the left fornix and the right and left optic radiations. The bundles with the highest percentages of associated fibers are shown in Table 5, with an anatomical representation provided in Figure 4.

**Table 5:** Results of the anatomical bundle identification regarding inverse associations between FA and NfL plasmatic levels (FDR < 0.05, n=118). For a description of the table headings, see Table 2. See Supplementary Material S4 for the complete report of the identified bundles.

| **Bundle** | **Bundle voxel n** | **Bundle voxel int** | **%** |
|---|---|---|---|
| Corpus callosum tapetum | 44230 | 1788 | 4.042 |
| Corpus callosum forceps major | 117410 | 1669 | 1.421 |
| Corpus callosum forceps minor | 115178 | 1205 | 1.046 |
| Right thalamic radiation posterior | 35747 | 367 | 1.026 |
| **Spearman rho (all tracts) = -0.198 (p = 0.031), IC [2.5 – 97.5] = -0.374, -0.011, Adjusted β = -0.196** |  |  |  |

**Figure 4:**
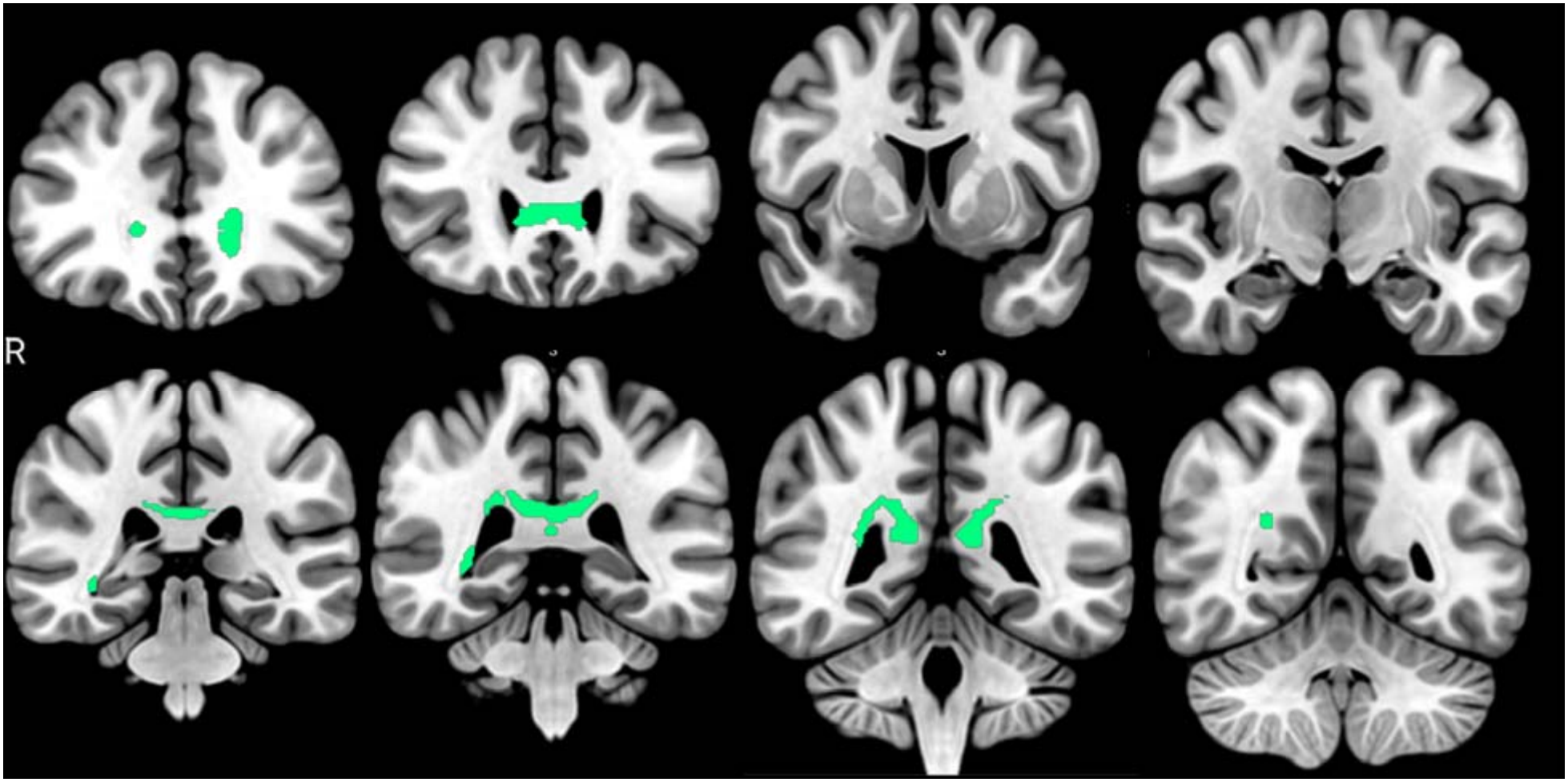
White matter tracts showing inverse associations between FA and plasma NfL levels (correlational tractography; Spearman partial correlation controlling for age and education; T-score threshold = 2.5; FDR < 0.05). Associations are concentrated in callosal fibers, with corpus callosum tapetum (4.04%), forceps major (1.42%), and forceps minor (1.05%) showing the highest proportions of compromised fibers. The complete list of identified tracts, including direct associations in limbi pathways, is provided in Table 6 and Supplementary Material S4.

Regarding direct associations, the cingulum and tapetum exhibited the strongest associations. A complete report of these findings, along with their anatomical localization, is presented in the Supplementary

**Table 6:** comparison of biomarker – FA correlation coefficients between pre and post WMH burden correction. Correlations with Aβ42/Aβ40 ratio are not reported, given that the reduction in sample size led to substantially lower variability in the biomarker, which generated the need for the corrections to be performed using only Aβ42 levels. For this, these results are merely exploratory.

| **Biomarker** | **Pre WMH correction** |  | **Post WMH correction** |  |
|---|---|---|---|---|
|  | **Spearman rho** | **p** | **Spearman rho** | **p** |
|---|---|---|---|---|
| **p-tau217 (inverse)** | -0.288 | 0.001 | Disappeared |  |
| **p-tau217 (direct)** | 0.174 | 0.051 | Disappeared |  |
| **GFAP (inverse)** | -0.274 | 0.002 | -0.297 | 0.001 |
| **GFAP (direct)** | 0.293 | 0.001 | 0.223 | 0.020 |
| **NfL (inverse)** | -0.198 | 0.031 | -0.259 | 0.006 |
| **NfL (direct)** | 0.220 | 0.017 | 0.155 | 0.107 |

### Associations of FA with NfL

Inverse associations with NfL were mainly localized in the corpus callosum. The right thalamic radiation was also identified, although with a markedly lower number of voxels involved. The quantification of these associations is reported in Table 6, with their anatomical representation shown in Figure 5.

**Figure 5:**
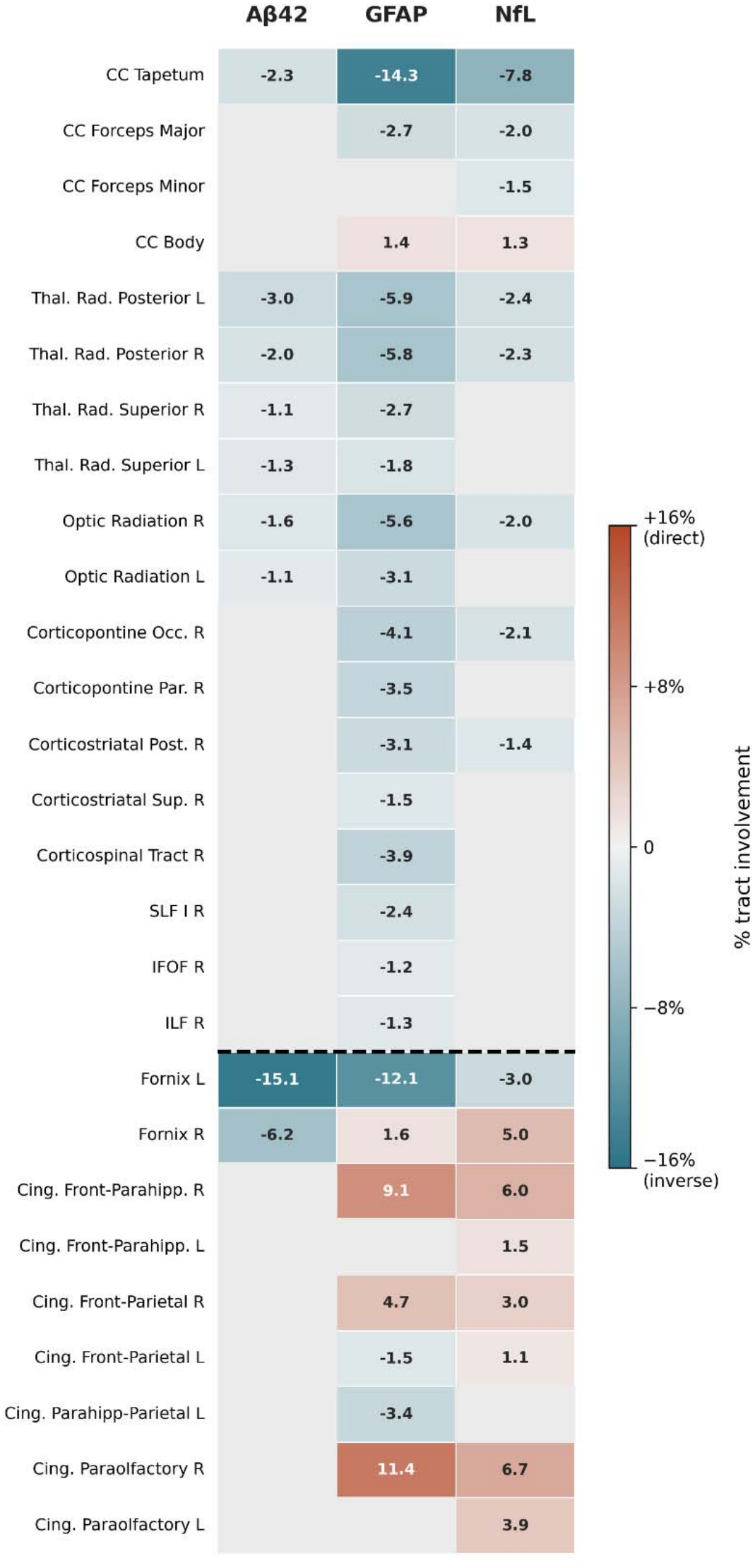
Heatmap showing the directionality and magnitude of FA associations for each plasma biomarker in the WMH-adjusted subsample (n = 124; Spearman partial correlation controlling for age, education, and TIV-normalized WMH volume; T-score threshold = 2.5; FDR < 0.05). Blue cells indicate inverse associations (higher biomarker levels associated with lower FA); red cells indicate direct associations (higher biomarker levels associated with higher FA). Numbers represent the percentage of tract volume with significant association relative to the HCP-1065 atlas definition. p-tau217 showed no significant associations in this subsample. Aβ42 was used as the amyloid-related variable due to insufficient variability in the Aβ42/Aβ40 ratio in this subsample. Legend: CC= Corpus callosum, SLF= Superior longitudinal fasciculus, IFOF= Inferior Fronto-Occipital Fasciculus, IFL = Inferior Longitudinal Fasciculus.

Finally, direct associations between NfL and FA were found in limbic circuit bundles, including the bilateral cingulum and fornix, as well as the corpus callosum tapetum. The quantification and anatomical representation of these associations are provided in the Supplementary Material S4.

To characterize hemispheric symmetry in our sample, we compared FA between 36 homologous left-right tract pairs using paired tests selected according to the normality of the paired differences (Shapiro-Wilk), with FDR-BH correction. Of the 36 tract pairs, 19 showed significant asymmetry (see Supplementary Material S5), with the largest effects observed in the parahippocampal cingulum, cerebellum, and superior corticostriatal tract. The direction of asymmetry in the arcuate fasciculus (left > right) was consistent with previously reported leftward lateralization of this tract in healthy populations (Takao et al., 2011). Because the left-handed subgroup was small (n = 3), we additionally repeated this analysis restricted to right-handed participants (n = 121); 35 of the 36 tract pairs retained the same significance status, indicating that the observed pattern of asymmetry is not attributable to the few non-right-handed participants in the sample.

### Associations between FA and AT(N) plasma biomarkers, after controlling by WMHs

When participants with Fazekas scores > 2 or without a score were excluded (n = 11) and TIV-normalized WMH volume was added as an additional covariate (n = 124), the pattern of associations between plasma biomarkers and FA changed. Given that the reduction in sample size led to substantially lower variability in the Aβ42/Aβ40 ratio, amyloid-related associations were assessed using plasma Aβ42 as the independent variable. Under this approach, inverse associations between Aβ42 and FA were identified, but it did not reach significance (Spearman rho = -0.125; p = 0.193, IC 2.5 – 97.5% = -0.317, 0.062. Adjusted β = - 0.074). No direct associations between Aβ42 and FA were observed in this subsample. Furthermore, after controlling for WMH burden, the associations between p-tau217 and FA disappeared.

For GFAP, inverse associations with FA persisted and was enforced, compared with the associations performed without WMH correction (Spearman rho = -0.297; p = 0.001, IC 2.5 – 97.5% = -0.456 , -0.115. Adjusted β = -0.251), with the highest proportions of affected fibers in the corpus callosum tapetum (14.33%) and left fornix (12.14%), followed by the bilateral posterior thalamic radiations (left: 5.86%; right: 5.84%), right optic radiation (5.64%), right corticopontine tract (occipital, 4.14%), right corticospinal tract (3.90%), right corticopontine tract (parietal, 3.51%), left cingulum (parahippocampal-parietal, 3.37%), left optic radiation (3.09%), and right corticostriatal tract (posterior, 3.06%). Direct association between GFAP and FA persisted, but was attenuated after WMH burden control (Spearman rho = 0.223; p = 0.020, IC 2.5 – 97.5% = 0.052, 0.378. Adjusted β = 0.252). These associations were identified in limbic-related tracts, primarily the right cingulum (paraolfactory, 11.37%), right cingulum (frontal-parahippocampal, 9.12%), right cingulum (frontal-parietal, 4.71%), and bilateral fornix (right: 1.55%; left: 1.04%).

For NfL, inverse association with FA persisted and was enforced with WMH correction (Spearman rho = - 0.259; p = 0.006, IC 2.5 – 97.5% = -0.436, -0.061. Adjusted β = -0.228), and were identified in callosal and posterior projection tracts, with the highest proportions of affected fibers in the corpus callosum tapetum (7.76%), corpus callosum forceps major (2.02%), corpus callosum forceps minor (1.50%), bilateral posterior thalamic radiations (left: 2.39%; right: 2.34%), and right optic radiation (1.99%). Direct associations between NfL and FA were attenuated and lost significance with the correction (Spearman rho = 0.155; p = 0.107, IC 2.5 – 97.5% = -0.015, 0.336. Adjusted β = 0.214).

No significant differences were found in biomarker levels between the individuals included and excluded from the analysis, based on their Fazekas score (see Supplementary Material S6).

A comparison of correlation coefficients between the pre- and post-WMH correction is presented in Table 7, and Supplementary Material S2. The directionality and proportion of affected tracts for each biomarker in this subsample are summarized in Figure 6. A complete anatomical report of this subsample analysis is available in Supplementary Material S7.

In the sensitivity analysis restricted to participants with SCC (i.e., without objective cognitive impairment), the pattern of compromised fibers was generally similar to that observed in the full cohort, suggesting that the observed associations were not exclusively driven by participants with objective cognitive impairment (see Supplementary Material S8 and S9).

## Discussion

The main findings of our study are that AT(N) plasma biomarkers were associated with FA in periventricular, callosal, and limbic tracts in a community-dwelling, non-demented cohort with cognitive complaints (GERO cohort). GFAP and p-tau217 showed the most extensive associations, whereas NfL showed a more restricted pattern, consistent with its specificity for axonal damage. Importantly, after excluding participants with Fazekas scores >2 (n = 124) and incorporating TIV-normalized WMH volume as an additional covariate, the associations with GFAP and NfL were maintained or even extended, whereas those with p-tau217 disappeared.

Considering participants’ cognitive characteristics, neuropsychological performance was heterogeneous across cognitive domains. Importantly, participant classification relied on the overall cognitive and functional profile rather than on isolated test performance. This approach is consistent with diagnostic frameworks in which a diagnosis of dementia requires cognitive impairment that interferes with independence in everyday activities, rather than low performance on a single cognitive measure alone (McKhann et al., 2011).

The differences observed between the full-sample analysis and the sensitivity analysis excluding participants with Fazekas scores >2 are in line with growing evidence indicating that WMHs reflect not only cerebrovascular injury but also neurodegenerative processes (Garnier-Crussard et al., 2023). Although WMHs have historically been treated as a marker of small-vessel disease (Pantoni, 2010), converging neuroimaging and post-mortem evidence indicates that WMHs reflect a mixed pathological substrate, combining small-vessel damage with amyloid- and tau-related processes, especially in periventricular and posterior territories (Alban et al., 2023; Du et al., 2024; Graff-Radford et al., 2019; McAleese et al., 2021). Lower CSF Aβ42 has been specifically associated with WMH burden in the splenium of the corpus callosum and posterior thalamic radiations (Alban et al., 2023), regions that overlap almost exactly with the territories driving the p-tau217 and Aβ42/Aβ40 associations observed in our primary analysis. In addition, longitudinal work has shown that WMH and amyloid burden interact synergistically to accelerate cognitive decline, with tau accumulation partially mediating this interrelationship (Du et al., 2024).

From this perspective, excluding high-Fazekas subjects and including TIV-normalized WMH volume as a covariate would not isolate the purely neurodegenerative share of FA variance. Rather, it would remove a composite source of variance encompassing both vascular damage and a portion of the amyloid- and tau-related microstructural damage that co-localizes with WMH-prone tissue. As a result, this statistical control would remove not only vascular-related FA variance but also FA variance driven by AT(N) pathology, potentially masking its associations with plasma biomarkers. Together, these statistical and biological considerations offer an evidence-based explanation for the divergent associations observed across biomarkers in the sensitivity analysis.

For GFAP and NfL, the pattern of FA associations extends well beyond periventricular regions into projection and association fibers (e.g., corticospinal, corticopontine, cingulum), where WMH burden is typically low and vascular and amyloid pathologies overlap less (Dang et al., 2026; Feng et al., 2025). The dominant variance in FA in these regions is therefore unlikely to be driven by WMHs, so adjusting for WMH burden and excluding subjects with the highest vascular load reduces confounding and reveals a more anatomically specific pattern of associations with astroglial and axonal injury markers. This interpretation is consistent with evidence that FA reductions precede the development of overt WMHs and partially mediate age- and amyloid-related WMH accumulation (Sun et al., 2025), positioning GFAP and NfL as plasma markers of an upstream microstructural process. In contrast, FA associations involving p-tau217, as well as the direct Aβ42 association, were topographically concentrated in posterior periventricular tracts (the tapetum, optic radiations, posterior thalamic radiations, and posterior corticostriatal fibers), precisely the territories in which DTI-derived microstructural changes and macroscopic WMHs exhibit the greatest spatial overlap. Thus, our interpretation of these results is that when WMH burden is statistically removed in this cross-sectional setting, a disproportionate share of the variance driving these associations would also be removed, not because the underlying biology is absent, but because it cannot be statistically separated from the co-localized vascular contribution, given the available sample size and design.

Biological evidence further supports this statistical interpretation. Reactive astrogliosis, indexed by GFAP, has been proposed to represent an early astrocytic response to white matter vulnerability that precedes overt lesion formation (Bettcher et al., 2024; Pereira et al., 2021). The expansion of GFAP–FA associations into projection systems that have not yet developed macroscopically visible damage is consistent with this model, in which plasma GFAP would capture an upstream stage of white matter degeneration before microstructural alterations progress to FLAIR-detectable WMHs. Longitudinal studies are needed to test this hypothesis directly.

The disappearance of p-tau217–FA associations after WMH adjustment is, in our view, the most striking finding of the secondary analysis and deserves careful interpretation. First, the loss of these associations should not be construed as evidence against a relationship between p-tau217 and white matter microstructure. p-tau217 is among the most specific plasma markers of ADD pathology, showing strong concordance with tau-PET findings even at preclinical and prodromal stages and across diverse populations (Caviedes et al., 2026; Kim et al., 2025; Orellana et al., 2026; Palmqvist et al., 2020). In our primary analysis, virtually all inverse p-tau217 associations were concentrated in posterior periventricular tracts (corpus callosum tapetum, optic radiations, posterior thalamic radiations, fornix, and posterior corticostriatal fibers), the same anatomical territory in which WMH-related amyloid and tau pathology preferentially accumulates (Parent et al., 2026). This topographic profile differs sharply from that of GFAP, whose associations extend into projection and association tracts that are less anatomically proximal to periventricular lesion-prone zones. Under these conditions, we propose that two mechanisms, which are not mutually exclusive, are likely to operate. First, because the variance in FA within posterior periventricular regions simultaneously reflects vascular small-vessel pathology and ADD-related (amyloid- and tau-related) microstructural injury, controlling for WMH volume removes a portion of the ADD-pathology-driven variance together with the vascular contribution. p-tau217, being closely linked to ADD pathology, is therefore disproportionately affected. This is consistent with evidence showing that medial-temporal tau accumulation partially mediates the synergistic effect of WMH and amyloid on cognitive decline (Du et al., 2025). Second, the robust direct p-tau217 associations in the right cingulum and bilateral fornix observed in the primary analysis likewise did not persist. Given that the same limbic direct associations were partially preserved for GFAP and NfL in the WMH-adjusted subsample, and that the p-tau217 subsample is smaller (n = 115 after exclusion for missing data), reduced statistical power may offer an additional explanation.

Importantly, this interpretation does not imply that p-tau217 is unrelated to white matter microstructure. Rather, it suggests that its associations with white matter are topographically restricted to regions where, in cross-sectional analyses with moderate sample sizes, they cannot yet be reliably distinguished from vascular contributions. As tau pathology spreads beyond these territories with advancing disease stage, p-tau217–FA associations are likely to become more spatially extensive and less susceptible to WMH adjustment. Longitudinal studies are necessary to adequately test this hypothesis.

A separate but related issue was the methodological need to switch from the ratio to Aβ42 alone in the filtered subsample. We justify this choice on two complementary grounds: first, the Aβ42/Aβ40 ratio in the n = 124 subsample showed insufficient between-subject variability for the connectometry algorithm to yield a stable correlational signal, with most values clustering below the commonly used range of 0.045– 0.047 (Ashton et al., 2024; Palmqvist et al., 2024). This pattern restricts the ratio’s analytic utility in a population-stratified, WMH-adjusted subsample. Second, using Aβ42 alone to examine its association with white matter microstructure is methodologically defensible, as previously reported by Bettcher et al. (2024), who used plasma Aβ42 in a sample spanning the cognitive spectrum to demonstrate synergistic effects of GFAP and amyloid on fornix FA, MD, and RD (Bettcher et al., 2024). The finding that emerged with this approach, only an inverse association between Aβ42 and FA in the bilateral fornix (15.1% of fibers affected on the left and 6.2% on the right), is biologically coherent. The fornix is among the earliest white matter pathways to show microstructural compromise along the AD continuum (Kantarci, 2014; Mielke et al., 2012), and is particularly susceptible to early amyloid-related neurodegeneration. The fact that the direct association between Aβ42 and FA in posterior periventricular tracts observed in the primary analysis disappeared after WMH adjustment, while the inverse fornix association was preserved, again fits the dual-pathology framework: the periventricular signal was confounded by WMH-related variance, whereas the limbic signal reflects early amyloid-related microstructural compromise not driven by macroscopic vascular damage.

A persistent challenge in interpreting these results is the absence of locally validated reference intervals for plasma biomarkers of neurodegeneration and glial reactivity in Chilean and broader Latin American populations (McGlinchey et al., 2024). To date, no reference ranges for Aβ42, Aβ40, p-tau217, GFAP, or NfL have been formally established in regional cohorts; clinical interpretation therefore currently relies on cut-off values derived primarily from North American and European cohorts. Despite this, we consider that our work not only demonstrates that biologically meaningful associations with white matter microstructure can be detected even without locally validated cut-offs, but also expands the empirical base from which future regional reference work can build. Given this, longitudinal, multimodal, and locally validated studies are crucial to establishing a more complete neurobiological interpretation of our results.

Several limitations should be acknowledged. First, men were underrepresented in the sample, a pattern commonly observed in aging research due to differential participation rates in health-related studies (Abdelnour et al., 2017; Forno et al., 2025), which limits the generalizability of the findings across sex. Future studies should take into account the evidence regarding differential associations between sex and AT(N) plasma biomarkers (Aksnes, 2025; Cheng et al., 2025; Coughlan et al., 2026; Labonte et al., 2025). Second, the absence of a cognitively unimpaired control group restricts the interpretability of the observed associations. Although the sensitivity analysis restricted to participants with SCC without objective cognitive impairment showed a similar pattern of compromised fibers, this subgroup cannot be considered equivalent to a cognitively unimpaired control group because all participants were recruited in the context of cognitive complaints. Likewise, the lack of a clinical dementia group prevents us from positioning these findings along the full continuum of cognitive decline. Future studies incorporating both CU participants and individuals at more advanced stages of impairment will be necessary to determine the stage-specificity of these associations. Third, although hand dominance was recorded and the sample was overwhelmingly right-handed (97.8%), the small number of left-handed participants (n = 3) precluded an adequately powered statistical comparison between handedness groups. A sensitivity analysis restricting the lateralization comparison to right-handed participants yielded materially unchanged results. Fourth, gold-standard biomarkers of AD pathology, such as CSF measures or amyloid PET, were not available due to limited accessibility and invasiveness, particularly in Global South settings (Daly et al., 2026; McGlinchey et al., 2024). Fifth, the cross-sectional design of the study does not permit inferences about temporal precedence between molecular and microstructural changes. The interpretive framework we propose, in which plasma GFAP and NfL may capture upstream microstructural processes preceding the emergence of WMHs, remains a hypothesis that must be tested longitudinally. Sixth, white matter bundle identification was based on probabilistic atlas-based labeling of reconstructed tracts (HCP-1065); while widely used, this approach introduces uncertainty related to inter-individual anatomical variability and partial overlap between adjacent fiber pathways, particularly in densely organized periventricular regions. Finally, the methodological switch from the Aβ42/Aβ40 ratio to Aβ42 alone in the WMH-adjusted subsample, although justified on statistical and biological grounds and supported by prior literature (Bettcher et al., 2024; Roseborough et al., 2017), introduces a non-trivial change in the operationalization of amyloid pathology between analyses; the results pertaining to Aβ42 should therefore be interpreted as complementary rather than directly comparable to those derived from the ratio in the primary analysis.

In conclusion, in this community-dwelling sample of Latin American older adults with cognitive complaints but without dementia, plasma biomarkers of neurodegeneration, astroglial reactivity, and axonal injury were associated with white matter microstructural alterations across a topographically coherent set of fiber bundles. The behavior of these associations under stringent control for macroscopic cerebrovascular pathology revealed a clear biological dissociation: associations of GFAP and NfL with FA were preserved and even spatially amplified, whereas associations of p-tau217 and the direct periventricular component of the Aβ42 signal were largely attenuated.

Interpreted in light of converging evidence that WMHs reflect mixed vascular and amyloid–tau pathology, especially in periventricular and posterior regions, these results suggest that plasma GFAP and, to a lesser extent, NfL are the most robust biomarkers associated with white matter microstructural vulnerability at the pre-dementia stage in this population, indexing astroglial and axonal processes biologically distinct from macroscopic small-vessel disease. p-tau217, in turn, captures microstructural change within a topographically constrained territory that remains statistically inseparable from co-occurring vascular pathology in moderate-sized cross-sectional designs, a finding that should not be read as evidence against its biological relevance, but rather as a limit on its detectability under WMH adjustment at this disease stage. The persistence of an inverse Aβ42–FA association in the fornix after WMH control highlights the limbic system as an early site of amyloid-related microstructural compromise, independent of vascular pathology.

By linking accessible blood-based markers to diffusion-derived measures of large-scale white matter integrity in a Latin American cohort with high cerebrovascular comorbidity, our work extends the empirical basis for globally representative interpretive frameworks for plasma biomarkers. Longitudinal, multimodal, and regionally anchored research, incorporating CU and clinical dementia groups, locally validated reference ranges, and integrated microstructural–macrostructural follow-up, will be essential to determine whether the microstructural signals captured here represent a transitional stage preceding the emergence of overt white matter lesions, and to refine the role of plasma GFAP, NfL, p-tau217, and Aβ42 as indicators of early white matter vulnerability across diverse global populations.

## Supporting information

Supplemental tables and script

## List of abbreviations

ADD: Alzheimer’s disease dementia
Aβ42: Amyloid-beta 42
Aβ40: Amyloid-beta 40
Aβ42/Aβ40: Amyloid-beta 42/40 ratio
CDR: Clinical Dementia Rating Scale
CU: Cognitively unimpaired
DSI Studio: Diffusion spectrum imaging studio
DTI: Diffusion tensor imaging
dMRI: Diffusion magnetic resonance imaging
FA: Fractional anisotropy
FDR: False discovery rate
FLAIR: Fluid-attenuated inversion recovery
GERO: Geroscience Center for Brain Health and Metabolism cohort
GFAP: Glial fibrillary acidic protein
HCP-1065: Human Connectome Project 1065-subject white matter atlas
ICBM-152: International Consortium for Brain Mapping 152-subject template
MCI: Mild cognitive impairment
MMSE: Mini-Mental State Examination
MNI: Montreal Neurological Institute
MRI: Magnetic resonance imaging
NfL: Neurofilament light chain
ODF: Orientation distribution function
PET: Positron emission tomography
PFAQ: Pfeffer Functional Activities Questionnaire
p-tau217: Phosphorylated tau at threonine 217
QSDR: Q-space diffeomorphic reconstruction
SCC: Subjective cognitive complaint
SD: Standard deviation
TE: Echo time
TIV: Total intracranial volume
TIP: Topology-informed pruning
TR: Repetition time
WMH: White matter hyperintensities

## Acknowledgments

This work was supported by ANID/FONDAP/15150012. PFR is supported by ANID Doctorate Scholarship Nº 21252107 / 2025. FH is supported by ANID Doctorate Scholarship Nº 21211340 / 2021 and ANID/FONDAP/15150012. VM is supported by ANID/FONDECYT de Iniciación 11251578, ANID/FONDECYT Exploración 13240170 and AARG-25-1490306 from the Alzheimer’s Association. JCC is supported by ANID/FONDECYT #1240807. FAC is supported by ANID/FONDAP/15150012, Michael J. Fox Foundation for Parkinson’s Research (17303) and Financiamiento Basal para Centros Científicos y Tecnológicos de Excelencia Centro Ciencia & Vida (FB210008). CDA is supported by ANID/FONDECYT Regular (1210622 & 1250091), ANID/NAM22I0007, ANID/PIA/ANILLOS ACT210096, Alzheimer’s Association (AARGD-24-1310017), ANID/FOVI240065, ANID/Proyecto Exploracion 13240170 and Multi-partner consortium to expand dementia research in Latin America [ReDLat, supported by Fogarty International Center (FIC), National Institutes of Health, National Institutes of Aging (R01s AG075775, AG057234, AG082056 and AG083799, CARDS-NIH75N95022C00031), Alzheimer’s Association (SG-20-725707), Rainwater Charitable Foundation – The Bluefield project to cure FTD, and Global Brain Health Institute)]. PG is supported by ANID AC3E CIA250006. AS is supported by ANID/FONDECYT/1231839; ANID/Proyectos de Exploración 13220082, Multi-partner consortium to expand dementia research in Latin America [ReDLat, supported by Fogarty International Center (FIC), National Institutes of Health, National Institutes of Aging (R01s AG075775, AG057234, AG082056 and AG083799, CARDS-NIH75N95022C00031), Alzheimer’s Association (SG-20-725707), Rainwater Charitable Foundation – The Bluefield project to cure FTD, and Global Brain Health Institute)], and Alzheimer’s Association GBHI ALZ UK-20-639295.

The authors would like to give special thanks to the patients and the team of the Memory and Neuropsychiatry Clinic, Hospital del Salvador - Faculty of Medicine, Universidad de Chile.

## Conflict of Interest Statement

The authors declare no conflicts of interest.

## Data availability statement

The data that support the findings of this study are available from the corresponding author upon reasonable request. Access to data from the GERO cohort is subject to ethical approval by the Scientific Ethics Committee of the Eastern Metropolitan Health Service in Santiago, Chile, and requires a formal data-sharing agreement with the research team. Requests for data access should be directed to the corresponding author.

## Supplementary material provided

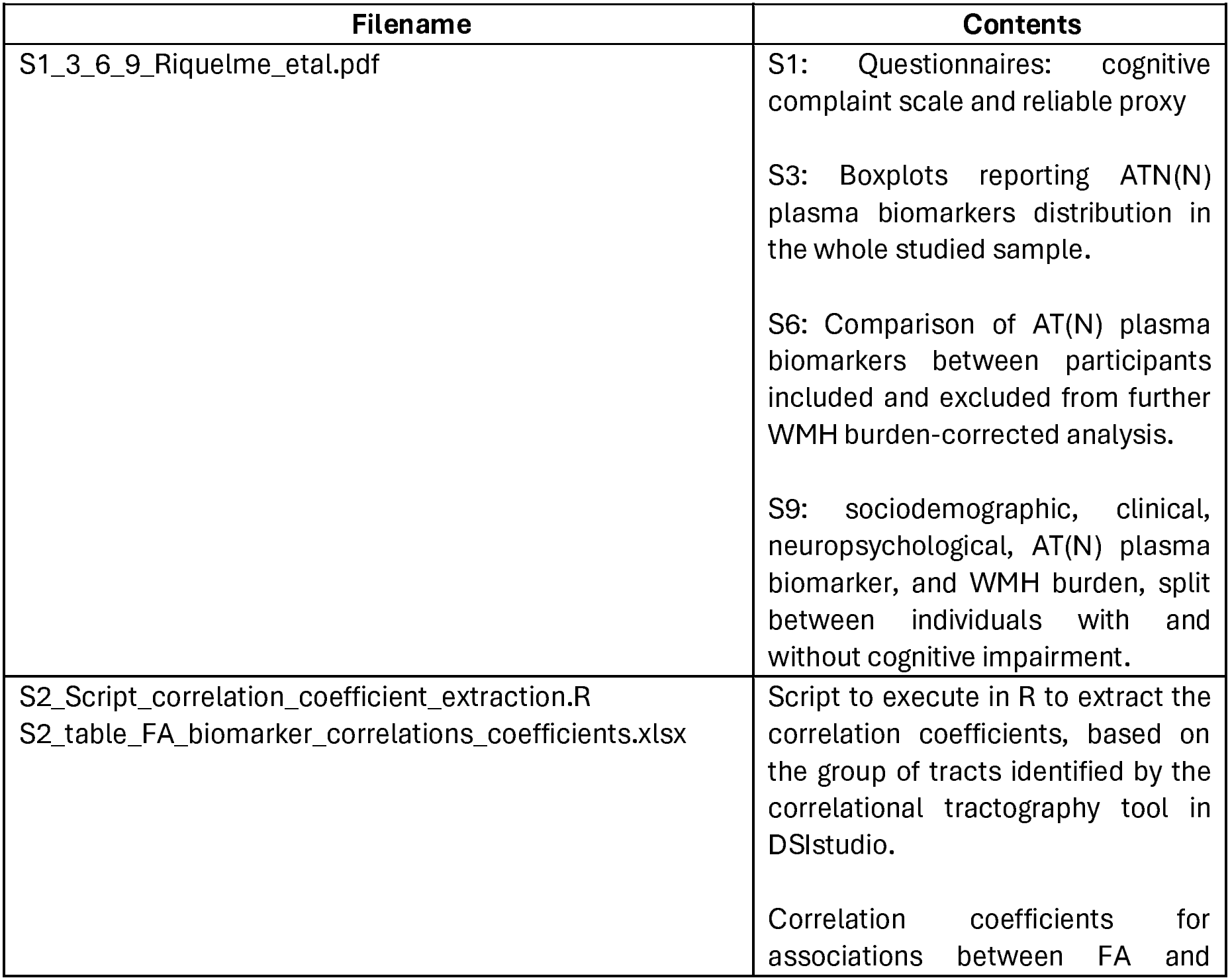
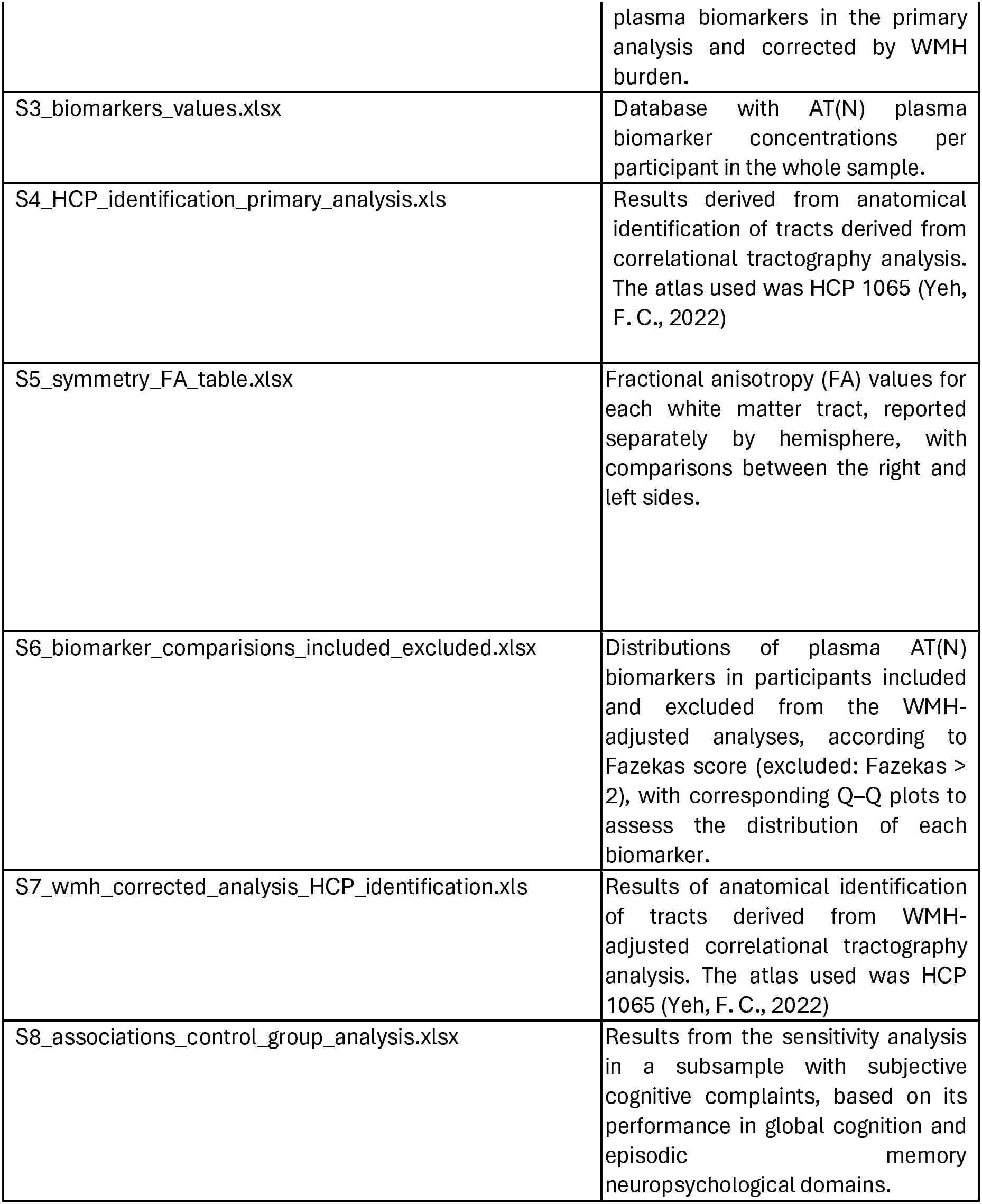

## Notes

### Competing Interest Statement

The authors have declared no competing interest.

