## Supplemental documents for "Associations between Fractional Anisotropy in White Matter Bundles and AT(N) Plasma Biomarkers in Individuals with Cognitive Complaints without dementia"

1. Department of Medical Technology. Faculty of Medicine, Universidad de Chile. Chile.
2. Geroscience Center for Brain Health and Metabolism (GERO). Santiago, Chile.
3. Neuropsychology and Clinical Neuroscience Laboratory (LANNEC). Interdisciplinary Nucleus for Physiology, Biophysics and Pathophysiology. Institute of Biomedical Sciences. Faculty of Medicine. Universidad de Chile. Chile.
4. PhD in Sciences / Neuroscience Program. Faculty of Sciences. Universidad de Valparaíso. Chile.
5. School of Medical Technology. Faculty of Medicine. Universidad de Chile. Chile.
6. Neurobiology of Integrative Systems (NoIS Lab). Facultad de Ciencias Biológicas. Pontificia Universidad Católica de Chile. Chile.
7. Department of Psychiatry, University of Pittsburgh, Pittsburgh, PA 15213, USA
8. Latin American Brain Health Institute (BrainLat), Universidad Adolfo Ibanez, Santiago, Chile.
9. Memory and Neuropsychiatric Center (CMYN) Hospital del Salvador & Faculty of Medicine, Universidad de Chile
10. Center for Integrative Biology, Faculty of Sciences, Universidad Mayor, Santiago 8580745, Chile.
11. Department of Chemistry and Biochemistry, University of California Santa Barbara, Santa Barbara, CA, 93106, USA.
12. Department of Neuroscience and Physiology, Upstate Medical University, NY, 13210, USA
13. Buck Institute for Research on Aging, Novato, CA, USA.
14. Center for Aging Research and Healthy Longevity, Faculty of Sciences, Universidad Mayor, Santiago, Chile.
15. Centro Científico y Tecnológico de Excelencia Ciencia & Vida, Fundación Ciencia & Vida, Santiago, Chile
16. Global Brain Health Institute (GBHI), Trinity College Dublin (TCD), Dublin, Ireland.
17. Department of Electrical Engineering. Faculty of Engineering. Universidad de Concepción. Chile
18. East Department of Neurological Sciences. Faculty of Medicine. Universidad de Chile. Chile.
19. Department of Neurosciences. Faculty of Medicine. Universidad de Chile. Chile. 7500922
20. Neurology and Psychiatry Department, Clínica Alemana – Universidad del Desarrollo. Santiago, Chile.

**Corresponding author**:
Andrea Slachevsky.
Geroscience Center for Brain Health and Metabolism (GERO). Santiago, Chile
Av. Salvador 486. Providencia. Santiago, Chile.

https://orcid.org/0000-0001-6285-3189

**Supplementary material S1:** Questionnaires of cognitive complaints referred to the participants and their proxy

**Cognitive Complaint Scale – Participant questionnaire**

1. **How is your memory compared to the way it was 10 years ago?**
    - Better
    - Same
    - Worse

2. **Do you remember things less well than you did a year ago?**
    - Yes
    - No
    - I do not know

3. **How is your ability to remember accurately the names of close friends and relatives?**
    - Very good
    - Good
    - Bad
    - Very bad

    **If in 3, the responses were Bad or Very Bad → <u>Positive cognitive complaint participant</u>.**

    **If no → <u>Continue</u>**

4. **Do you believe your memory has become worse?**
    - Yes
    - No
    - I do not know

5. **Do you usually forget where things are?**
    - Yes
    - No
    - I do not know

6. **Do you have difficulties finding the right words during a conversation?**
    - Yes
    - No
    - I do not know

7. **Do you remember accurately appointments?**
    - Very good
    - Good
    - Bad
    - Very bad

8. **Do you have trouble remembering things that have recently happened?**
   - Yes
   - No
   - I do not know

9. **Do you ever forget what you were told yesterday or the day before?**
   - Yes
   - No
   - I do not know

**If questions 8 and 9 were responded as "Yes" → Positive Cognitive Complaint Participant.**

**If no → Negative Cognitive Complaint Participant.**

10. **Do you ever start doing something and forget what you were doing?**
    - Yes
    - No
    - I do not know

11. **Do you ever go to a room to look for something and forget what you came for?**
    - Yes
    - No
    - I do not know

**Questions from**: Rabin LA, Smart CM, Crane PK, et.al. (2015) *"Subjective Cognitive Decline in Older Adults: An Overview of Self-Report Measures Used Across 19 International Research Studies"*. J Alzheimers Dis. Sep 24;48. doi: 10.3233/JAD-150154.

**Instrument validated in**: Molina Molina-Donoso M, Parrao T, Meillon C, et.al. (2023). *"Assessing subjective cognitive decline in older adults attending primary health care centers: what question should be asked?"* J Clin Exp Neuropsychol. May;45(3):313-320. doi: 10.1080/13803395.2023.2221399.

**Cognitive Complaint Scale – Reliable Proxy questionnaire**

**Proxy initial evaluation**

**Inclusion questions**

1. **Have you, or has someone close to you, noticed significant changes in memory of your relative that cause you concern?**
    - Yes
    - No

2. **Are these memory changes concerning enough for you to consider seeking professional evaluation or consultation?**
    - Yes
    - No

3. **Have you sought, or has anyone suggested that you seek professional evaluation for this problem?**
    - Yes
    - No

    **If you have not sought evaluation, please indicate the reason(s):** *(Select all that apply.)*

    a) *Did not consider it necessary, so did nothing.*
    b) *Did not consider it necessary and used home remedies.*
    c) *Preferred to consult a pharmacist for medication related to the health problem.*
    d) *Preferred to consult a specialist in alternative medicine (e.g., vibrational medicine, biomagnetism, reiki, iridology, Bach ffower therapy, Eastern medicine, Indigenous medicine, naturopathy, or homeopathy).*
    e) *Considered seeking consultation, but did not have time.*
    f) *Considered seeking consultation, but did not have money.*
    g) *Considered seeking consultation, but found it difficult to reach the healthcare facility.*
    h) *Requested an appointment but did not obtain one.*
    i) *Obtained an appointment, but it has not yet taken place.*
    j) *Obtained an appointment, but did not attend*
    k) *Don't know / Do not apply*

**AD8**. Below are a series of situations that refer to the person you are reporting on. In your case, Mr./Ms. (participant's name). Please answer “Yes” IF YOU CONSIDER THAT THERE HAS BEEN ANY CHANGE in the aspect indicated over the last few years, due to cognitive problems (thinking, memory, etc.)

| **Nº** | **Situation** | **Yes** | **No** | **I do not know / Does not apply** |
|---|---|---|---|---|
| 1 | Problems making judgments and decisions (being deceived, making poor financial decisions, giving inappropriate gifts). | | | |
| 2 | Loss of interest in hobbies and activities (e.g., has stopped doing activities they used to enjoy) | | | |
| 3 | Repeat questions, comments, or things that are said. | | | |
| **4** | **Difficulty learning how to use tools, appliances, or devices (e.g., video or DVD player, computer, microwave, remote control, cell phone, or cordless phone) *** | | | |
| **5** | **Forget the correct month or year *** | | | |
| 6 | Difficulty managing complicated financial matters (e.g., making payments, paying taxes, bills, receipts, etc.) | | | |
| 7 | Difficulty remembering appointments and things you must do | | | |
| 8 | Problems with reasoning and/or memory are daily occurrences, not occasional ones. | | | |

**Proxy ABCD-Q**

| Nº | Ítem | Strongly disagree | Disagree | Agree | Strongly agree |
|---|---|---|---|---|---|
| 9 | His / Her memory or concentration difficulties make it harder for you to live independently. | | | | |
| 10 | He / She is more indifferent to the feelings or problems of others. | | | | |
| 11 | He / She is less aware when your behavior bothers or annoys others. | | | | |
| 12 | His / Her comments, or behavior have become more inappropriate (e.g., poor manners, use of profanity, or other socially inappropriate actions). | | | | |
| 13 | He / She finds it more difficult to organize and express his / her ideas. | | | | |
| 14 | He / She finds it harder to make appropriate decisions | | | | |
| 15 | He / She finds it harder maintaining concentration on what is doing. | | | | |
| 16 | He / She is confused about the order in which events occurred. | | | | |
| **17** | **He / She has become more forgetful*** | | | | |
| 18 | He / She tends to repeat him(her)self more often. | | | | |
| 19 | He / She has greater difficulty remembering conversations that took place a few days ago | | | | |

If one of the questions in bold was answered as "Yes" (for questions 4 or 5), or Agree / Strongly Agree (for question 17) → **Positive Cognitive Complaint according to the proxy.**

**If NO → Negative Cognitive Complaint according to the proxy.**

**References on validation of cognitive complaint questions**

Slachevsky, A., Parrao, T., Lillo, P., Forno, G., Villagra, R., Thumala, D., Amieva, H., Zitko, P., Galvez, R., Assar, R. and Wenk, E. (2017), [P2–443]: SUBJECTIVE MEMORY COMPLAINTS AND COGNITIVE DECLINE. Alzheimer's & Dementia, 13: P806-P806. https://doi.org/10.1016/j.jalz.2017.06.1099

Parrao, T., Thumala, D., Lillo, P., Forno, G., Amieva, H., Wenk, E., Toledo, C. and Slachevsky, A. (2018), P3-333: SUBJECTIVE MEMORY COMPLAINTS AND COGNITIVE PERFORMANCE IN A POPULATION OF HEALTHY OLDER ADULTS ATTENDING PRIMARY CARE CENTERS IN SANTIAGO, CHILE. Alzheimer's & Dementia, 14: P1208-P1208. https://doi.org/10.1016/j.jalz.2018.06.1694

**Supplementary material S3:** Boxplots of AT(N) plasma biomarker distributions in the whole sample (n = 135). In the boxplots, the central line represents the median value, the colored box represents the interquartile range, the upper whisker represents the maximum (Q3 + 1.5*IQR), and the lower whisker represents the minimum (Q1 – 1.5*IQR).

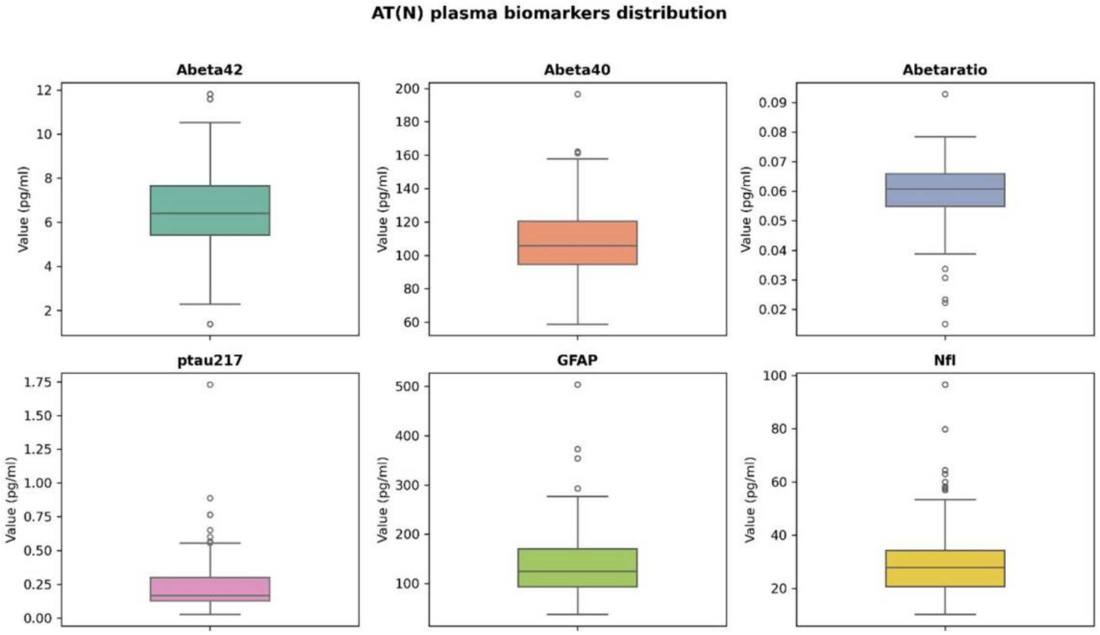

**Supplementary material S6:** Distributions of plasma AT(N) biomarkers in participants included and excluded from the WMH-adjusted analyses, according to Fazekas score (excluded: Fazekas > 2), with corresponding Q–Q plots to assess the distribution of each biomarker. In boxplots, the orange line represents the median value, box represents the interquartile range, upper whisker represents maximum (Q3 + 1.5*IQR), and lower whisker represents minimum (Q1 – 1.5*IQR). p-values reported correspond to comparison statistic test used in each variable, (*) means Mann-Whitney U test, and (**) means t-test.

## Aβ40

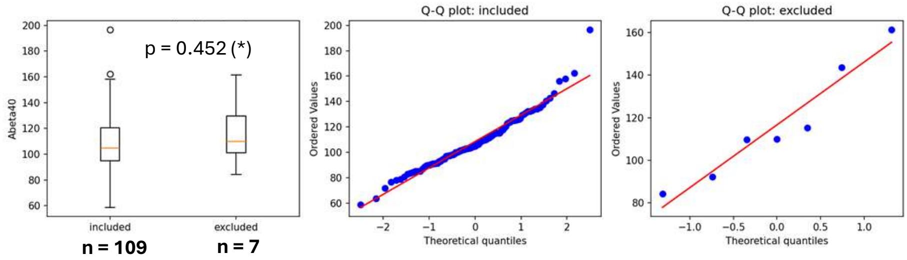


## Aβ42

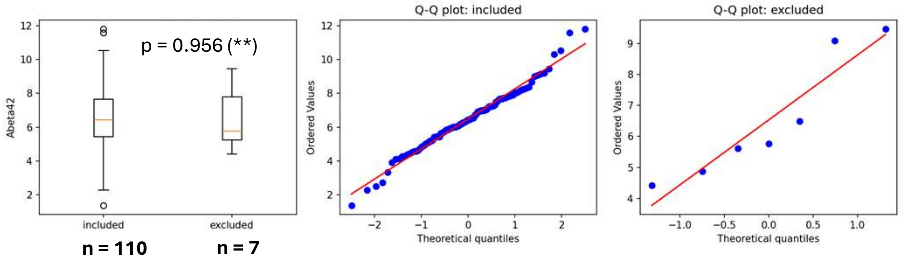


## Aβ42/40

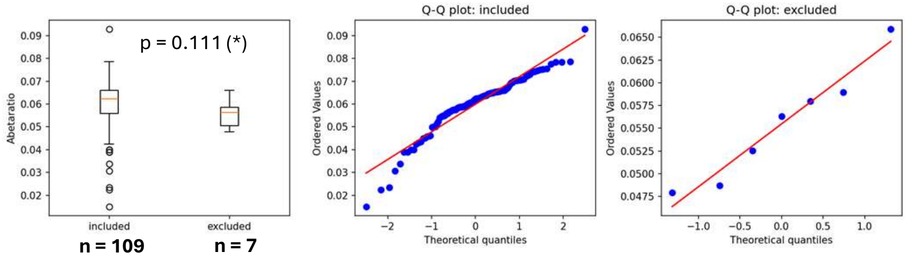


### p-tau 217

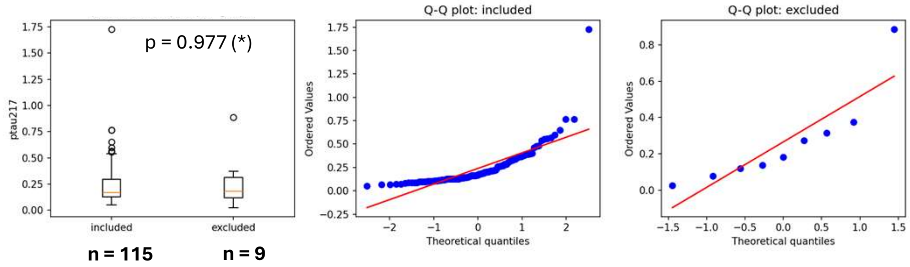


### GFAP

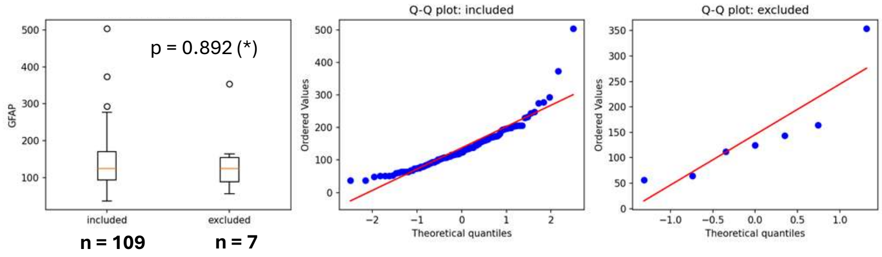


### Nfl

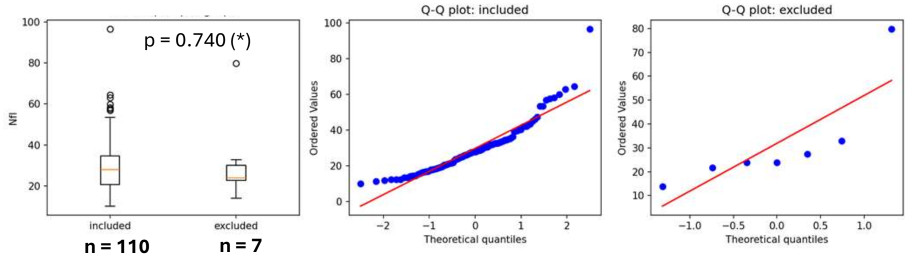

**Supplementary material S9:** sociodemographic, clinical, neuropsychological, AT(N) plasma biomarker, and WMH burden, split between individuals with and without cognitive impairment.

Values are mean ± SD; median [min, max], unless otherwise indicated.

| Variable | Total sample | No cognitive impairment | With cognitive impairment |
|---|---|---|---|
| *n* | *135* | *88* | *47* |
| ***Sociodemographics*** | | | |
| Sex, M : F, n (%) | 27 : 108 (20 : 80) | 21 : 67 (24 : 76) | 6 : 41 (13 : 87) |
| Age (years) | 76.09 ± 4.93; 75 [69, 92] | 75.73 ± 4.81; 74.5 [69, 92] | 76.79 ± 5.14; 75 [69, 89] |
| Education (years) | 9.36 ± 4.66; 8 [0, 24] | 10.55 ± 4.72; 10 [0, 24] | 7.15 ± 3.68; 6 [0, 17] |
| ***Clinical and global cognition*** | | | |
| MMSE total score | 27.09 ± 2.28; 28 [20, 30] | 28.01 ± 1.60; 28 [22, 30] | 25.35 ± 2.38; 26 [20, 30] |
| MoCA total score | 21.53 ± 4.38; 22 [11, 29] | 24.17 ± 2.52; 24 [13, 29] | 16.54 ± 2.31; 17 [11, 20] |
| CDR total score | 0.45 ± 0.16; 0.5 [0, 0.5] | 0.45 ± 0.16; 0.5 [0, 0.5] | 0.44 ± 0.16; 0.5 [0, 0.5] |
| PFAQ total score | 0.76 ± 1.28; 0 [0, 8] | 0.74 ± 1.39; 0 [0, 8] | 0.80 ± 1.07; 0.5 [0, 5] |
| ***Multi-domain neuropsychological assessment*** | | | |
| Semantic memory: FCSRT total recall | 47.31 ± 1.62; 48 [34, 48] | 47.49 ± 1.02; 48 [43, 48] | 46.95 ± 2.36; 48 [34, 48] |
| Episodic memory: FCSRT delayed free recall | 11.67 ± 2.66; 12 [3, 16] | 12.11 ± 2.48; 12 [5, 16] | 10.84 ± 2.84; 11 [3, 16] |
| Attention: CTT A (sec) | 98.44 ± 53.83; 83 [24, 360] | 86.44 ± 42.93; 77 [24, 283] | 123.95 ± 65.30; 114.5 [54, 360] |
| Executive function: CTT B (sec) | 234.12 ± 134.16; 194 [59, 540] | 186.58 ± 100.58; 156 [59, 540] | 335.15 ± 141.57; 336 [106, 540] |
| Language: SydBat Naming | 24.74 ± 3.27; 25 [13, 30] | 25.90 ± 2.48; 26 [19, 30] | 22.45 ± 3.46; 23 [13, 28] |
| ***Plasma biomarkers and WMH burden*** | | | |
| Aβ42 (pg/ml) | 6.48 ± 1.77; 6.40 [1.38, 11.81] | 6.57 ± 1.86; 6.52 [1.38, 11.81] | 6.32 ± 1.61; 6.10 [2.51, 9.47] |
| Aβ40 (pg/ml) | 108.96 ± 21.29; 105.87 [58.75, 196.61] | 107.47 ± 23.35; 102.85 [58.75, 196.61] | 111.64 ± 16.96; 109.84 [78.06, 143.64] |
| Aβ42/Aβ40 ratio | 0.05 ± 0.01; 0.060 [0.015, 0.092] | 0.061 ± 0.012; 0.063 [0.015, 0.093] | 0.057 ± 0.012; 0.059 [0.022, 0.078] |
| p-tau217 (pg/ml) | 0.23 ± 0.20; 0.167 [0.025, 1.728] | 0.23 ± 0.16; 0.167 [0.066, 0.887] | 0.26 ± 0.28; 0.186 [0.026, 1.729] |
| NfL (pg/ml) | 29.91 ± 14.01; 27.88 [10.13, 93.61] | 29.88 ± 15.32; 27.07 [10.13, 96.61] | 29.98 ± 11.47; 30.44 [11.90, 58.27] |
| GFAP (pg/ml) | 138.50 ± 70.80; 124.79 [36.98, 503.72] | 133.02 ± 60.97; 124.35 [37.38, 373.10] | 148.30 ± 85.55; 131.88 [36.98, 503.73] |

| Variable | Total sample | No cognitive impairment | With cognitive impairment |
|---|---|---|---|
| Total WMH (TIV normalized) | 0.00 ± 0.008; 0.008 [$8.16 \times 10^{-4}$, 0.051] | 0.010 ± 0.008; 0.008 [$1.08 \times 10^{-3}$, 0.052] | 0.010 ± 0.009; 0.007 [$8.17 \times 10^{-4}$, 0.052] |

Abbreviations: MMSE, Mini-Mental State Examination; MoCA, Montreal Cognitive Assessment; CDR, Clinical Dementia Rating; PFAQ, Pfeffer Functional Activities Questionnaire; FCSRT, Free and Cued Selective Reminding Test; CTT, Color Trails Test; SydBat, Sydney Language Battery; NfL, neurofilament light chain; GFAP, glial fibrillary acidic protein; WMH, white matter hyperintensities; TIV, total intracranial volume.
